# Coopting the Lap system of *Pseudomonas fluorescens* to reversibly customize bacterial cell surfaces

**DOI:** 10.1101/358705

**Authors:** T. Jarrod Smith, Holger Sondermann, George A. O’Toole

**Affiliations:** Department of Microbiology and Immunology, Geisel School of Medicine at Dartmouth Rm 202 Remsen Building, Hanover, NH 03755; Department of Molecular Medicine, College of Veterinary Medicine, Cornell University, Ithaca, New York 14853

**Author notes:** Address correspondence to George A.O‘Toole.

**Keywords:** biofilm, cell surface, engineering, T1SS, adhesin

## Abstract

Initial attachment to a surface is a key and highly regulated step in biofilm formation. In this study, we present a platform for reversibly functionalizing bacterial cell surfaces, with an emphasis on designing biofilms. We engineered the Lap system of *Pseudomonas fluorescens* Pf0-1, which is normally used to regulate initial cell surface attachment, to display various protein cargo at the bacterial cell surface and control extracellular release of the cargo in response to changing levels of the second messenger cdi-GMP. To accomplish this goal, we fused the protein cargo between the N-terminal retention module and C-terminal secretion signal of LapA, and controlled surface localization of the cargo with natural signals known to stimulate or deplete c-di-GMP levels in *P. fluorescens* Pf0-1. We show this system can tolerate large cargo in excess of 500 amino acids, direct *P. fluorescens* Pf0-1 to surfaces it does not typically colonize, and program this microbe to sequester the toxic medal cadmium.

---

The bacterial biofilm lifestyle is profoundly consequential to human health and industry. Although the concerning link between biofilm formation and increased antibiotic tolerance has been known for some time (1, 2), only recently have the benefits of some surface-attached communities become appreciated. Such beneficial roles include competitively excluding pathogen colonization (3), bioelectricity generation (4), and enhancing bioleaching (5, 6). Furthermore, recent microbiome research cataloging various beneficial relationships between bacterial biofilms and their human host has caused a for therapeutic purposes (7). One goal of synthetic biology is to program bacterial cell surfaces to perform customized functions under an exclusive set of environmental conditions, such as binding a defined surface or remediating a toxic metal from an environment.

The first stage of biofilm formation is when a bacterium makes initial contact with a substratum while later stages are focused on reinforcing the biofilm matrix after committing to a surface. To establish a biofilm, many bacteria employ surface-associated adhesins to initially bind a surface (8–10) and subsequently secrete adhesins, small amyloid proteins and/or complex exopolysaccharides to “glue” the bacteria together within the biofilm (11, 12). Synthetic biologists have exploited strategies for manipulating these two stages of biofilm formation; however, many of these biofilm engineering tools are restricted to displaying relatively small protein domains. To promote initial contact, bacterial surfaces have been modified to display surface tags (13), light-responsive amphiphiles (14), photoswitchable proteins (15), and photoswitchable azobenzene linkers (16). Likewise the small, self-assembling amyloid protein CsgA of *E. coli* has been functionalized for nanoparticle templating as well as mercury bioremediation (17, 18). However, these strategies are often unable to accommodate large domains (>60 aa), limiting their versatility and thus downstream applications. Interestingly, exploiting the natural bacterial decision-making process to tune initial attachment and biofilm formation has also been largely overlooked, with researchers favoring synthetic, UV light– or blue light-oriented strategies.

Here, we describe a new approach for reversibly customizing the bacterial cell surface using the Lap system of *P. fluorescens* Pf0-1, which is naturally used to promote initial attachment and thus biofilm formation by a variety of microbes (19). The Lap system was recently characterized as a novel subgroup of T1SS transporters and substrates (20). This system is comprised of 3 components: a type 1 secretion system (T1SS) apparatus (LapEBC), a giant adhesin (LapA), and a inside-out regulatory component (LapGD) that controls levels of LapA at the cell surface in response to the second messenger c-di-GMP (Figure 1). LapA is a ~520 kDa adhesin with extensive internal repeats that are sandwiched between an N-terminal retention module and C-terminal secretion signal (20, 21). The level of surface-associated LapA corresponds with cellular c-di-GMP concentrations, allowing rapid, tunable changes in biofilm assembly and disassembly (22). The inner membrane-bound c-di-GMP receptor, LapD, controls the activity of the periplasmic LapA-targeting protease, LapG. LapG cleaves LapA at a characterized dialanine site; however, when bound to c-di-GMP, LapD sequesters LapG to protect LapA, and thereby promote LapA surface localization and thus biofilm formation (23, 24).

**Figure 1.**
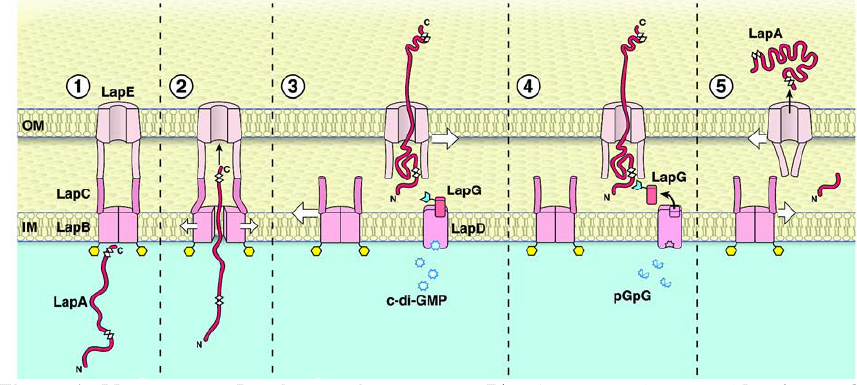
Model of the LapA adhesion system. The C-terminal domain of LapA includes the secretion signal, which engages the T1SS (panel 1). LapA is secreted C-terminal end first (panel 2), and when c-di-GMP levels are high the N-terminal retention domain anchors LapA to the cell surface via retention in the outer membrane component of the T1SS (panel 3). If c-di-GMP levels fall, the LapG protease is released from the LapD receptor and LapG is free to cleave the N-terminal retention domain of LapA (panel 4). The LapA lacking the N-terminal domain is release from the cell surface (panel). A portion of this figure was published previously (20). Figure copyright William Scavone, 2018. Used with permission.

To develop the Lap system as a platform for customizing the *P. fluorescens* Pf0-1 cell surface, we sought to first delete the gene encoding the giant adhesin, *lapA*, then use the regions critical for LapA cell surface localization (20) and secretion (21) to deliver and control cell surface release of different cargo proteins of interest. We have previously shown a 3XHA epitope tag N-terminally fused to LapA‘s C-terminal secretion signal (pC235, 5012M-5246S; Figure 2A, B) is secreted directly into the supernatant independent of LapG activity (20). To determine if LapA‘s N-terminal domain (1M-272I) is sufficient to display the 3XHA-tagged secretion signal of LapA at the cell surface, we fused this N-terminal region to C235 to generate the pN272 construct (pN272; Figure 2A). The pC235 and pN272constructs were then expressed in the *lapA* and *lapAlapG* mutant backgrounds and assayed for cell surface localization and LapG-dependent release into the extracellular environment. Here, the lapA mutant has a functional LapG protease capable of cleaving the N-terminal retention module of LapA (see Figure 1) while the lapAlapG mutant lacks the protease and thus the retention module of LapA remains intact and functional. We predicted the N272 fusion protein, despite containing only ~10% of the full-length LapA protein, should localize to the cell surface and require LapG proteolysis for release into the supernatant similarly to the full-length LapA.

**Figure 2.**
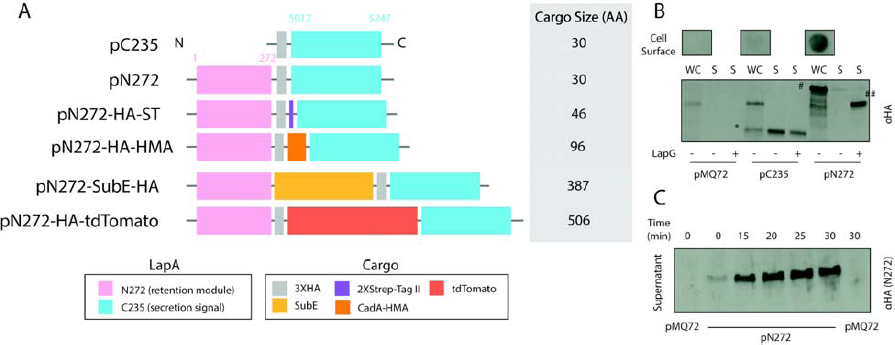
(A) Schematic representation of N272 variants described in this study. The legend indicates the various domains in each construct. The C-terminal domain of LapA (light blue) contains the secretion signal, and is common to all constructs. The pink domain is the “retention module” of LapA and is required to anchor LapA to the cell surface (see Figure 1). (B) Cell surface association and regulation of the N272 construct requires the N-terminal retention module. Cell surface, whole cell (WC), and supernatant (S) levels of the indicated proteins after subculturing for 6 hr. The presence (+) of absence (-) of LapG in the strain is indicated. (C) Controlled extracellular release of surface associated N272 in response to phosphate starvation and removal of the calcium chelator. Strains were subcultured for 5.5 hr in a LapG-inhibiting medium (0 min), the medium was replaced with a LapG-activating medium and the supernatant sampled over 30 min for the presence of cleaved N272.

Western blot analysis indicated the N272 fusion is displayed at the cell surface (Figure 2B, cell surface). Furthermore, consistent with previous studies with full-length LapA (25), extracellular release of N272 requires LapG proteolysis, as indicated by comparing the molecular weight of N272 in the whole cell (WC, intact N272, #) and supernatant fractions (S, proteolyzed N272, ##) when LapG is absent (-) or present (+) (Figure 2B, bottom, far right). Conversely, control strains expressing LapA‘s 3XHA-tagged secretion signal demonstrate this variant lacking LapA‘s retention module is unable to associate with the surface and is secreted directly into the extracellular environment independent of LapG activity (Figure 2B, p235, middle, *).

Cell surface levels of LapA can be tuned by modulating cellular levels of c-di-GMP or by inhibiting the proteolytic activity of the calcium-dependent protease, LapG. Phosphate robustly stimulates c-di-GMP production in *P. fluorescens* Pf0-1 while phosphate-limiting conditions activate the Pho regulon, leading to transcriptional activation of the phosphodiesterase RapA and depletion of c-di-GMP levels, thus decreasing LapA at the cell surface and reducing biofilm formation (26). Alternatively, LapG, a calcium-dependent as EGTA or citrate, leading to LapA retention and biofilm formation independently of c-di-GMP (27). We took advantage of this knowledge to determine if we could control release of the pN272 construct into the supernatant. The lapA mutant expressing pN272 was grown in a high phosphate medium supplemented with the calcium chelator 0.2% citrate to discourage LapG proteolysis. This LapG-inhibiting medium was then exchanged with the same base medium, except depleted for phosphate and lacking citrate, both of which stimulate LapG activity. Cleaved N272 in the supernatant fraction was then monitored for 30 minutes. Western blot analysis indicates LapG activation enriches the supernatant fraction with cleaved N272 peptide within 15 minutes, illustrating the rapid responsiveness of this system (Figure 2C).

Given that LapA naturally contains an extensive and complex domain architecture, we hypothesized this LapA-based platform could be utilized to reversibly display various protein cargo on the bacterial cell surface. To test this idea, we cloned several cargo proteins into the N272 system, as shown in Figure 2A. We then assayed for LapG-dependent, cell-surface release of the cargo to determine if this platform could be applied to differentially functionalize the *P. fluorescens* Pf0-1 cell surface. The spectrum of cargo tested ranged from a cytoplasmic Heavy-Metal Associated domain (the HMA from the ABC transporter CadA of *Listeria*), to a protease secreted by a Gram-positive bacterium (subtilisin E of *Bacillus subtilis*), as well as the fluorescent protein tdTomato and small epitope tags (3XHA and 2X Strep-tactin) (Figure 2A). Notably, all of the cargo tested was displayed at the cell surface and released in response to LapG activity (Figure 3); however Western analysis of the whole cell fraction indicated some variability in cargo stability displayed and release from the *P. fluorescens* Pf0-1 cell surface suggests this system can tolerate large, multifunctional cargo and can display proteins and domains of cytoplasmic or extracellular origin.

**Figure 3.**
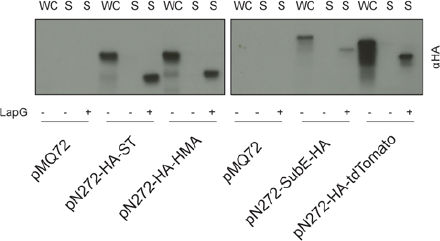
Extracellular release of cargo requires LapG activity. Western blot analysis of the whole cell (WC) and supernatant (S) fractions from the *lapA* mutant (LapG, +) and the *lapAlapG* mutant (LapG, -) mutant strains expressing the indicated constructs.

Because the *lapA* mutant does not form a biofilm under our laboratory conditions, we next asked if a cargo displayed at the cell surface in the pN272 variants could direct *P. fluorescens* Pf0-1 to bind a surface of interest. To test this idea, we performed a competitive binding assay with *lapA* mutants expressing either empty vector or pN272-SubE-HA mixed at a 1:1 ratio. The cell mixture (input) was applied to protein G magnetic beads bound to αHA anti-body to determine if presentation of the HA epitope conferred selective binding to the functionalized beads. After a short incubation period, the resin-bound bacteria were isolated from the mixture with a magnet and the free-floating population was collected and characterized (Figure 3, right). While the input contained equal numbers of binding to non-binding cells, the output, which represents cells that could not bind the functionalized beads, was almost exclusively binding-defective cells expressing the empty vector (Figure 4, right). These data suggest the Lap system may be utilized to direct *P. fluorescens* to functionalized or novel surfaces for various biotechnological applications.

**Figure 4.**
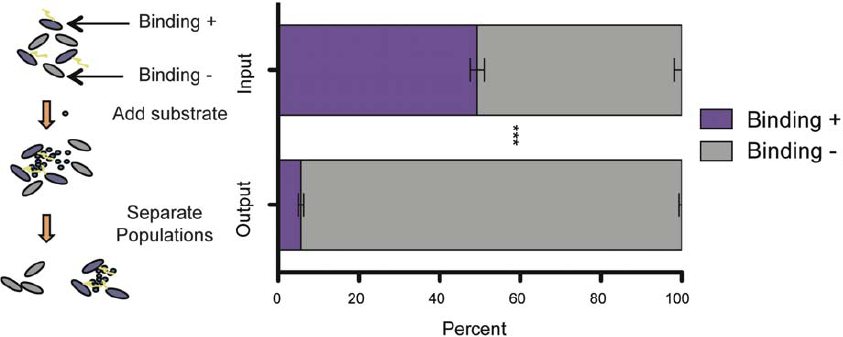
Competitive binding assay (outlined on left) between *P. fluorescens* Pf0-1 *lapA* mutants expressing pN272-SubE-HA (binding-positive) or empty vector (binding-negative). The two strains were mixed at a 1:1 ratio (input) then incubated with αHA antibody-bound protein G magnetic resin. Cells bound to the resin were removed from the mixture using a magnet, and the supernatant fraction with unbound cells collected (output). Cells from the input and output were plated. Colony PCR was performed on 100 random colonies from the input and output to enumerate cells carrying empty vector (pMQ72) or pN272-SubE-HA. Error bars are SEM of three biological replicates. Two-way ANOVA statistical analysis was performed (***, p<0.0001).

**Figure 5.**
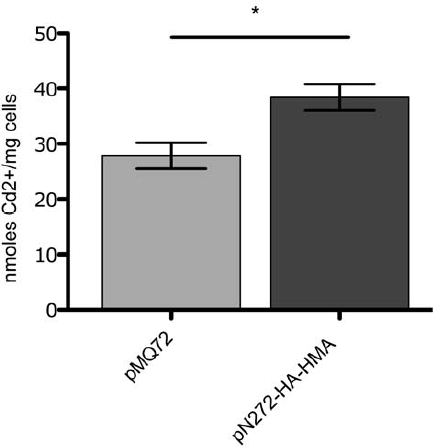
*P. fluorescens* Pf0-1 displaying the HMA domain from CadA of L. monocytogenes show increased cadmium binding. The *P. fluorescens* Pf0-1 *lapA*lapG mutant expressing the indicated plasmids were subcultured for 5.5 hr and then exposed to 10 μM Cadmium sulfate for 30 minutes. Weighed cell pellets were resuspended in equal volume of buffer and lysed. Cadmium levels were determined using ICP-MS. Error bars are SEM of three biological replicates. Two-tailed t-test was performed (*, p<0.05).

Designing microbes for bioremediation purposes is also of immense interest to synthetic biologists. Thus, we next wanted to ask if the Lap system could be used to design *P. fluorescens* Pf0-1, a natural plant symbiont, to bind the heavy-metal cadmium, which is highly toxic to plants. To test this idea, we expressed the cytoplasmic cadmium-binding HMA domain from the P1-type ATPase CadA of compare cellular cadmium levels between the *P. fluorescens* Pf0-1 *lapAlapG* mutant *L. monocytogenes* (28) in our N272 system (Figure 2A, pN272-HA-HMA) and assayed for cadmium binding. We used ICP-MS to expressing pN272-HA-HMA or empty vector (pMQ72) after being exposed to 12 μM cadmium sulfate (CdSO_4_) for 30 minutes. The modest, but statistically significant increase in bound cadmium suggests the cytoplasmic HMA domain is functional when displayed at the *P. fluorescens* Pf0-1 cell surface. These data are consistent with the HMA domain sequestering cadmium, suggesting the Lap system may be engineered for bioremediation purposes.

In summary, we present a platform to customizing bacterial cell surfaces using the Lap system from *P. fluorescens* Pf0-1 and demonstrate its usefulness in biofilm design and bioremediation. Like most T1SS, the Lap system can accommodate large protein cargo unsuitable for other cell-surface display platforms, expanding potential downstream applications of this system. The customized cargo displayed at the cell surface can be tuned by modulating levels of the secondary messenger c-di-GMP or through chemical inhibition of the calcium-dependent protease LapG, allowing rapid, controlled biofilm assembly and disassembly. Together, these features make the Lap system an attractive platform for functionalizing the bacterial cell surface. Although we have demonstrated this proof of concept in *P. fluorescens* Pf0-1, various Gram-negative bacteria encode the T1SS (19) suggesting it may be optimized to reversibly functionalize the cell surface of various Gram-negative bacteria.

## Materials and Methods

### Plasmids, Bacterial Strains, and Growth Conditions

The plasmid pMQ72 (29) was used as the backbone for all the constructs engineered for this study. The N-terminus and C-CadA-HMA domain from *L. monocytogenes* DNA sequences were ordered from IDT. The gene coding for subtilisin E was cloned from Bacillus subtilis 168. The pRSF-Duet plasmid was a gift from Prof. Holger Sondermann. Plasmid carrying tdTomato was a gift from Prof. Deb Hogan. S17 E. coli was purchased from Life Technologies. The *P. fluorescens lapA* and *lapA*lapG clean deletion mutant strains, described previously (20), carrying pMQ72-based plasmid were grown overnight in LB + 30μg/mL Gentamycin and subcultured with rotation in K10T-1 (30) for 6 hours unless noted otherwise.

### Cloning of pN272 Cargo

Yeast cloning was used to fuse cargo with LapA N– and C-terminal elements into pMQ72. The N– and C– terminus of LapA was amplified using PCR primers designed with ends homologous to *SmaI* digested pMQ72 to orient insertion. Each cargo was amplified with PCR primers designed with ends homologous to either the 3‘ end of LapA‘s N-terminus or 5‘ end of LapA‘s C-terminus to orient insertion.

### Western Blot Analysis

Standard practices for Western blot analysis and cell-surface LapA detection were used to detect the 3XHA or 6XHIS epitopes engineered into the pN272-based constructs, as reported (21). For whole cell analysis, cells were normalized and resuspended in 1X SDS-page loading buffer. For supernatant analysis, the supernatants were concentrated in Amicon centrifugal 4 mL 30K NMWL spin columns (Millipore Cat. #UFC803096) and protein levels normalized following protein quantification using the Pierce BCA assay kit (Thermo #23227).

### Competitive Binding Assay

*p.fluorescens lapA* mutants were subcultured in K10T-1 +0.4% sodium citrate (Fisher, Cat. No. S279-500) to inhibit LapG activity, normalized, and applied to Pierce Protein G Magnetic Resin (Cat #88847) prepared and pre-incubated with 1 μg a-HA anti-body (BioLegend #901503) according to the manufacture‘s suggestions. Cells were incubated with anti-body bound magnetic resin at room temperature for 1 hr. The resin-bound fraction was separated with a magnet, and then the medium fraction containing the unbound bacteria was plated. Colony PCR was used to enumerate cells carrying pN272-SubE-HA and empty vector (pMQ72).

### Cadmium Binding

*P. fluorescens lapAlapG* mutants expressing pN272-HA-HMA or empty vector (pMQ72) were subcultured for 5.5 hours and exposed to 10μM Cadmium sulfate (Fisher Cat. # C19-500) for 30 min. To prepare for ICP-MS analysis, the dry cell pellets were weighed and resuspended in 25mM Tris pH7.4, then boiled for 20 min at 100°C. The cell debris was removed with centrifugation and the lysate submitted for analysis.

## Acknowledgement

This work was funded by supported by NIH grant R01GM123609 (to H.S. and G.A.O.). We thank M.L. Guerinot for assistance in designing the metal binding cargo. Metal analysis was performed by the Trace Elements Analysis Core, supported by NIH/NIEHS P42 ES007373 and NIH/NCI P30CA023108.

